# Characterization of widely conserved novel pigment production in *Bacillus subtilis* species

**DOI:** 10.1101/2024.11.28.625866

**Authors:** Rune Overlund Stannius, Christopher A. Dunlap, Estelle Morvan, Mélanie Berbon, Sophie Lecomte, Antoine Loquet, Ákos T. Kovács

## Abstract

*Bacillus subtilis* is widely studied in the microbial secondary metabolite (SM) field due to its rich variety of important natural products and genetic tractability. However, identification of novel SMs and their biosynthetic gene cluster (BGCs) has become increasingly difficult, especially in *Bacilli*, as the tools for screening and genome mining are dependent on clear function or similarity to already known BGCs. Pigments are SMs identified by their absorption of visible light, resulting in a certain color perceived by our eyes at sufficient concentrations. Thereby, pigments provide the evidence of a BGC without knowing the sequence or function. Expanding the known repertoire of SM BGCs with novel BGCs will further reinforce identification of a broader set of BGCs by mining tools such as antiSMASH. Here, we study a pigment observed in *B. subtilis* soil isolate MB9_B4 on certain media. We characterize the conditions where this pigment is produced and identify the corresponding BGC using a comparative genomic approach exploiting our strain collection containing other isolates with pigment production ability. The responsible BGC carried several genes, which were annotated as parts of the tryptophan biosynthesis pathway, possibly originating from a duplication and divergence of an originally primary metabolism. Identification of the pigment gene cluster additionally lead to the discovery of additional pigment BGC carrier *B. subtilis* isolates, some of which were described at the earliest in 1896 under the name *Bacillus aterrimus*, with a name referring to a dark pigmentation (the Latin “aterrimus” meaning very black). In addition, we employed solid-state nuclear magnetic resonance and Fourier transform infrared spectroscopies to characterize the chemical groups of the pigment. This study describes the chemical and biological features of a new class of SM BGC, which we hope will serve to improve the current BGC discovery pipelines in *Bacilli*.

## 1. INTRODUCTION

*Bacillus subtilis* is a Gram-positive soil dwelling bacterium that, due to its genetic tractability and spore development, has been one of the focal species in microbiology with decades of research into its lifecycle and sporulation (1–4). Concomitantly with sporulation, many *Bacilli* produce a wide array of pigments with varying structures and proposed roles, mainly as UV protection and reactive oxygen species scavenging benefitting spore resilience (5–9). However, pigments may have many roles beyond just their spectral properties and ability to scavenge reactive oxygen species. In *B. subtilis*, the most well studied pigment, pulcherriminic acid, has been shown to modulate iron availability by sequestering Fe(III) from competing organisms and preventing harmful Fenton reactions (10) and also influence biofilm development (11, 12). Pigments pose a promising target for study due to their intrinsic visibility without specific detection technology.

Pigments belong to the group of secondary metabolites (SMs) which are differentiated from primary metabolism by having specialized functions not directly related to the growth and reproduction of the producing organism. SMs are a rich source of novel therapeutics and functional molecules for society with a large majority of medicinal compounds either being SMs or derivatives of SMs (13, 14). However, discovery of truly novel SMs is becoming difficult, especially in the *Bacillus* genus; screening assays often remove SMs from their context and test for very specific functions, which further skews our knowledge towards well-known SM classes and functions while losing cryptically expressed SM biosynthetic gene clusters (BGCs) and restricting efforts to cultivable bacteria (15). One way to avoid reliance on expression and functional assays would be through sequence-based approaches, which can identify potential BGCs by similarity to already known clusters. Such approach has the added advantage of being culture independent, allowing access to the largely untapped diversity of microorganisms in nature (13). However, sequence-based approaches are only likely to find SM BGCs that are similar in some way to already defined BGC classes and therefore are unlikely to find truly novel SM BGCs (16).

Evolutionarily, SM biosynthetic genes are thought to originate from primary metabolism through duplication and diversification of enzymes in which the encoded proteins gain new substrate specificity or enzymatic action (17). More recently diverged BGCs are still highly similar to their progenitor genes from the primary metabolism and are often not identified by various BGC focused genome-mining tools as parts of the secondary metabolism. Alternatively, exploiting the evolutionary principles involved in the origin of BGCs has been proposed as a strategy with the genome-mining approach EvoMining, which circumvents aforementioned challenges by instead targeting enzyme family duplications and identifying potential BGCs by conserved genomic vicinity and phylogenetic reconstruction of the duplicated enzymes (16, 18). However, more examples of primary metabolism derived SMs are still valuable for the similarity-based methods and for better understanding the evolution of BGCs.

In this study, we identify a dark water-soluble pigment produced by a *B. subtilis* soil isolate, MB9_B4. Through a genome-wide association study (GWAS), we were able to identify the putative gene cluster associated with pigment production, which consisted of multiple genes annotated as homologues of the primary metabolism for tryptophan. Blast searches revealed several other isolates that carry the pigment BGC, some of which had been studied for their pigmentation in different media conditions under the name *Bacillus aterrimus* as early as in 1896 (19–21). The chemical nature of the pigment was assessed by high resolution solid-state nuclear magnetic resonance and Attenuated total reflection Fourier transform infrared (ATR-FTIR) spectroscopy. This study adds to the knowledge base of less well-studied SMs in *Bacilli* and provides an example of a primary metabolism-derived secondary metabolite, which could be useful for improving our BGC discovery pipelines.

## 2. MATERIALS AND METHODS

### 2.1 Strains, chemicals, and genetic modification

The list of strains used for the experiments can be found in Table S1, while additional isolates included in phylogeny are indicated in Table S1. Lysogeny broth (LB; Lennox, Carl Roth, 10 g/L tryptone, 5 g/L yeast extract and 5 g/L NaCl) was used for routine culturing. LBg (LB supplemented with 1% v/v glycerol), LBm (LB supplemented with 0.1 mmol/L MnCl_2_), LBgm (LB supplemented with both 1% v/v glycerol and 0.1 mmol/L MnCl_2_, based on (22)), MSgg (1.925 mmol/L KH_2_PO_4_, 3.075 mmol/L K_2_HPO_4_, adjusted to pH 7, 100 mmol/L 3-(N-morpholino)propanesulfonic acid (MOPS), 2 mmol/L MgCl_2_, 700 µmol/L CaCl_2_, 50 µmol/L MnCl_2_, 50 µmol/L FeCl_3_, 1 µmol/L ZnCl_2_, 2 µmol/L thiamine, 0.5% v/v glycerol, 0.5% w/v L-glutamic acid potassium salt, 50 µg/mL tryptophan, (23)), Potato dextrose agar (PDA, Carl Roth), and Kings B agar (KB, sigma Aldrich) were additionally used for assaying pigment production under various conditions. Antibiotics were used at the following final concentrations: Spectinomycin (spec) 100 µg/mL, kanamycin (kan) 5 µg/mL, tetracycline (tet) 10 µg/mL, chloramphenicol (chl) 10 µg/mL, erythromycin (erm) 5 µg/mL for erm^R^ strains and 1 µg/mL combined with 12.5 µg/mL lincomycin for MLS^R^ strains.

All genetic engineering was performed as described below using a modified version of the protocol described in (24). Briefly, overnight cultures were pelleted and resuspended in 100 µL MQ water. 10 µL of the concentrated cells was transferred into 2 ml competence medium (80 mmol/L K_2_HPO_4_, 38.2 mmol/L KH_2_PO_4_, 20 g/L glucose, 3 mmol/L Na_3_-citrate, 45 µmol/L ferric NH_4_-citrate, 1 g/L casein hydrolysate, 2 g/L K-glutamate, 0.335 µmol/L MgSO_4_·7H_2_O), which was put in a shaking incubator at 37 °C for 3.5 hours. Donor DNA was extracted from 1 ml of an overnight culture of the donor strain in LB using the Bacterial and Yeast Genomic DNA Purification Kit from EURx, which yielded a typical purified DNA concentration ranging from 50 – 150 ng/µL. 2 µL donor DNA was added to a fresh tube and mixed with 400 µL of competent cells and further incubated 2 hours at 37 °C before plating of 100 µL on LB agar plates with appropriate antibiotics for selection of transformants, and incubated overnight before selection of successful transformants.

### 2.2 Comparative genome analysis

To identify the gene cassettes correlating with pigment production, a pangenome was constructed using Panaroo (25–28) and subsequently used in a Genome-wide association study (GWAS) with pigmentation phenotypes in pyseer (29). Due to the low sample size of pigment producers (3 out of 13), hits were analyzed by hand focusing on co-localization of significant hits. Genomes were additionally re-annotated using bakta (30) at the default settings for any additional information on the identified genes.

### 2.3 Production of ^13^C,^15^N-labeled samples

The *B. subtilis* MB9_B4 GFP and MB9_B4 GFP Δ*yetJ*-PGC strains (Table S1) were cultured in 5 mL of LB, at 37°C, 120 rpm overnight, as pre-culture. 5 mL of labelled MSgg buffer (5 mM K_2_HPO_4_ / KH_2_PO_4_ pH7, 100 mM MOPS pH7, 0.5% L-glutamic acid ^13^C_5_ ^15^N, 0.5% glycerol ^13^C_3_, 50 μM tryptophane, 50 μM threonine, 2 mM MgCl_2_ 6H_2_O, 2 μM thiamine, 700 μM CaCl_2_, 100 μM MnCl_2_, 50 μM FeCl_3_, 1 μM ZnCl_2_) were then inoculated at 0.1% and cultured at 30°C without agitation for 5 days in order to trigger the bacillus biofilm at the surface. The media culture was collected with a syringe with a needle 21G, and centrifuged at 15 000 g, 30 min, 4°C in order to keep the supernatant. The pellet containing the biofilm was recovered with a spatula and used for spectroscopic analysis.

### 2.4 Solid-state NMR and Fourier transform infrared spectroscopies

^13^C,^15^N-labeled samples were analyzed by magic-angle spinning solid-state NMR. One dimensional ^13^C and ^15^N spectra were recorded on a 600 MHz Bruker magnet equipped with a 4 mm triple resonance probe. 11 kHz spinning frequency and high-power ^1^H decoupling (83 kHz) were used to record 1D INEPT using acquisition times of 20 and 15 ms for ^13^C and ^15^N spectra respectively. Chemical shifts were calibrated according to sodium trimethylsilylpropanesulfonate (DSS). ATR-FTIR spectroscopy was performed on a Nicolet iS50 FTIR spectrometer equipped with an MCT detector cooled with liquid N_2_ and an ATR accessory mounted with a germanium crystal (one reflection). Spectra were recorded with 100 scans and a spectral resolution of 2 cm^-1^.

## 3. RESULTS

### 3.1 Culture conditions for pigment production

When testing the antifungal activity of *B. subtilis* isolates on PDA medium (31), we observed that MB9_B4 produces a pigmented compound that diffuses into the agar medium. To get a better understanding of the conditions at which the pigment is produced, we screened pigmentation under a range of culture media (**Fig. 1**).

**Fig. 1.**
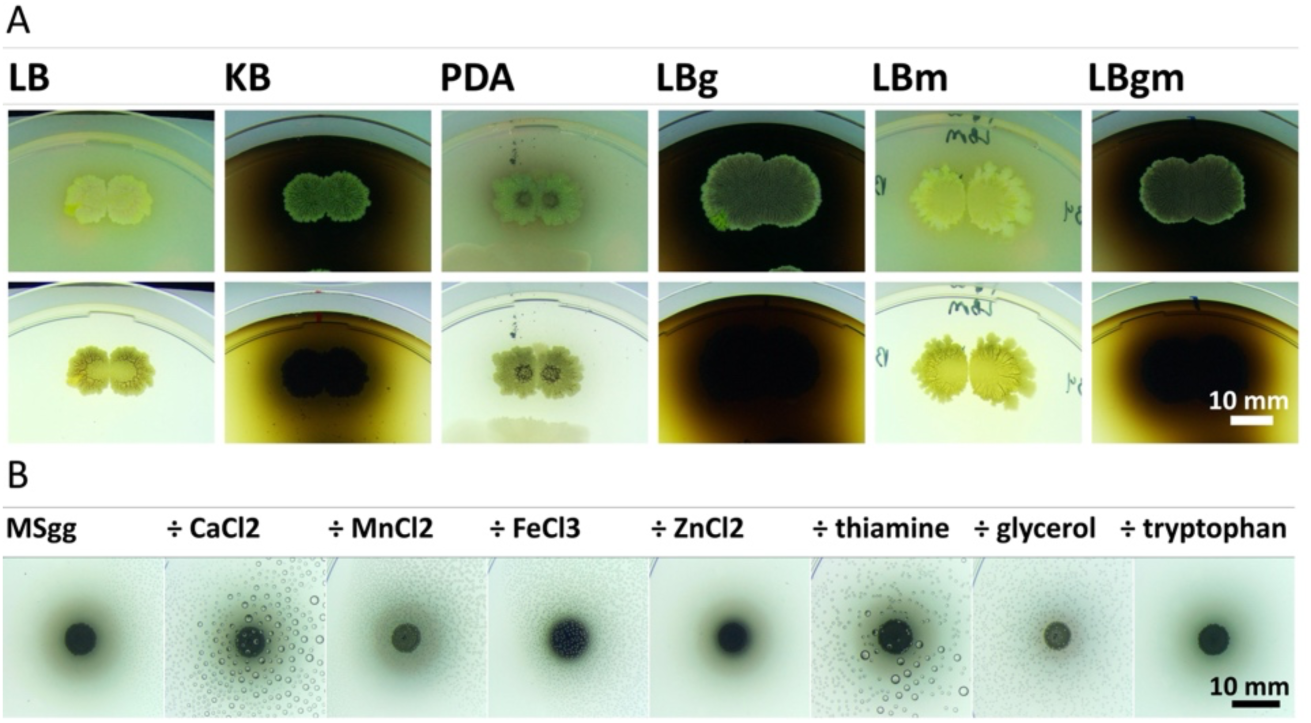
A) Pigmentation of MB9_B4 on various solid media after 5 days at room temperature imaged with either contrasting light from top (top row) or below (bottom row) the plate, scale bar corresponds to 10 mm. B) Medium component analysis of MB9_B5 pigmentation on MSgg agar after 5 days at room temperature, showing complete media to the left and each component omitted as noted above separate images with scale bar in the lower left corner corresponding to 10 mm.

In colony biofilm culturing, we were able to reproduce the same pigmentation observed in earlier studies on PDA (**Fig. 1A**). More interestingly, pigmentation was even stronger on biofilm inducing media such as MSgg (chemically defined biofilm-promoting medium) and LBgm (LB supplemented with glycerol and manganese that induces biofilm development in *B. subtilis*) compared to PDA, which suggested a possible involvement of the biofilm pathway or biofilm-inducing condition in pigment production. Further testing of LBg (LB supplemented with glycerol) and LBm (LB supplemented with manganese) featured pigmentation only on LBg, although it should be noted that LB itself contains trace amounts of manganese, which might be enough for induction when only glycerol is added. Lastly, pigmentation was also induced on King’s B agar, which is a known inducer of fluorescent pigments in *Pseudomonas*. Both MSgg, LBg, LBgm, and KB contains glycerol, which might indicate glycerol itself as a determinant factor and not biofilm induction through the glycerol-manganese biofilm inducing pathway noted in (22).

To probe the importance of certain additives, we performed a medium component analysis in MSgg agar, leaving out each component from the chemically defined medium, which revealed that pigment production did not depend on any single of the tested components; suggesting that glycerol-mediated induction of biofilm through the glycerol-manganese pathway is not the major inducer of pigment production (**Fig. 1B**).

Pigmentation was observed at 25 °C (although delayed by 1 day) and 35 °C, but was absent at 45 °C (**Fig. 2A**). We noted a variation in coloration, which was clearest on MSgg with low temperatures resulting in a more “bluish” pigment and higher temperatures leading to a “brownish” coloration. We tested if the variable coloration was a physiological response to the temperature or a chemical change in the pigmentation itself by sterile filtering diluted pigmented medium (which allowed easier evaluation of any color change) and incubating at higher temperatures, which resulted in a shift to a brown color indicating a temperature dependent decay (**Fig. 2B**).

**Fig. 2.**
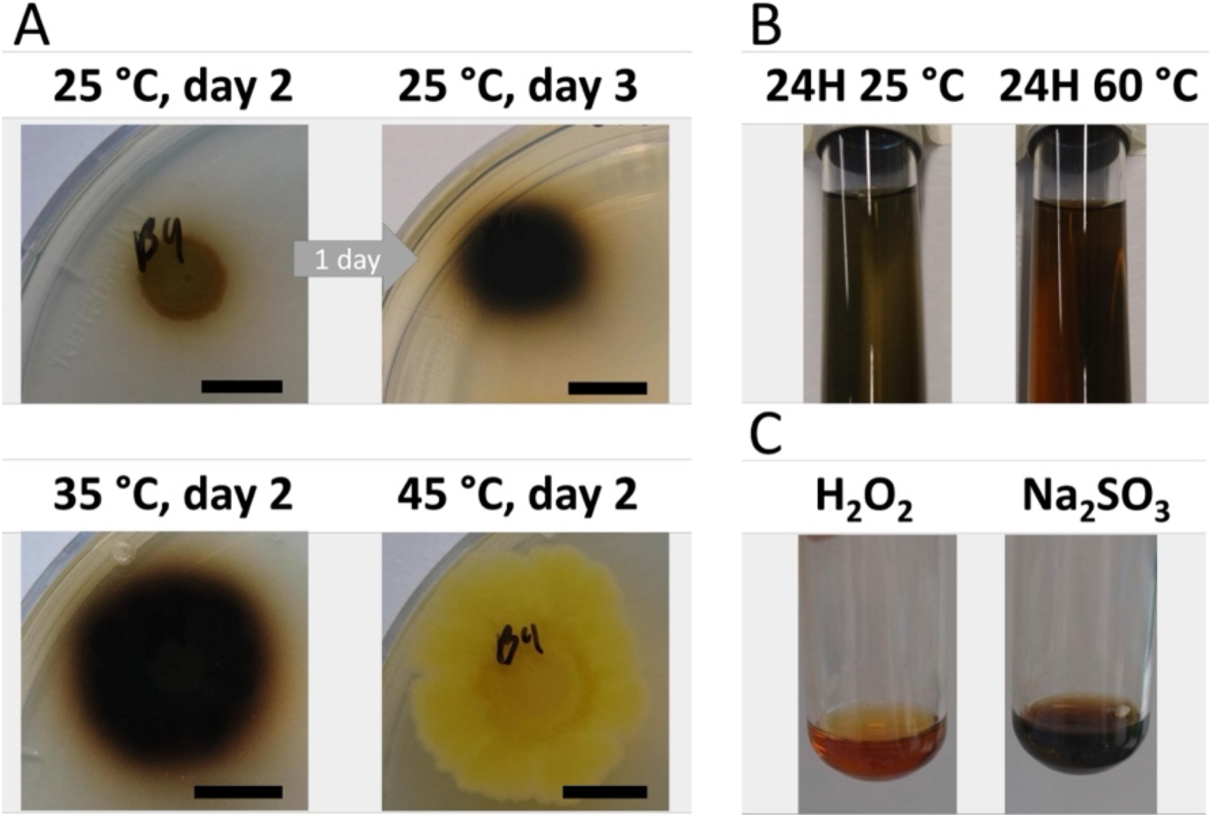
A) Pigmentation on MSgg agar plates kept for two days at 25, 35, and 45 °C, respectively, day 3 picture shown for 25 °C condition as pigmentation improved considerably, scale bar in each picture corresponds to 10 mm. B) Change in coloration of 10× diluted pigment kept for 24 hours at 60 °C compared to 25 °C. C) Effect of 24-hour treatment with 1% v/v H_2_O_2_ and 1% w/v Na_2_SO_3_.

To check if the change in coloration was due to oxidation, we further tested the effect of hydrogen peroxide (1% v/v H_2_O_2_) as well as the reducing agent sodium sulfite (1% w/v Na_2_SO_3_). Oxidizers were able to bleach the pigment while the reducing agent did seemingly not affect coloration (**Fig. 2C**). Additionally, the oxidization was irreversible; supplementation of reducing agent to the oxidized cultures resulted in no further change. It is possible that color variation is due to spectral mixing of other pigments and it is not caused by the change in the absorption spectra of the specific pigment, which might be perceived from such color change.

### 3.2 GWAS and confirmation by deletion

We screened 12 additional *B. subtilis* isolates from our strain collection (31) for pigmentation on MSgg agar plates to facilitate comparative genomics (**Fig. S1).** Here we found pigmentation in 3 out of the 12 isolates (MB9_B4, 63, and 75). Comparing the pigmented isolates with non-pigmented strains revealed the presence of genes in the pigmented strains that are annotated as homologues of the tryptophan biosynthetic operon co-located within a ∼16-kbp gene cluster between the *yetJ* and *yetK* loci, which we designated as PGC (pigment gene cluster) (**Fig. 3A**, **Table 1 and 2**). The connection between pigment production and the identified gene cluster was subsequently confirmed by deleting the *B. subtilis* MB9_B4 PGC between *yetJ* and *yetK* using homologous recombination of the upstream and downstream regions present in the *yetJ*::*kan^R^* and *yetK*::*kan^R^* mutant strains of the BKK library (BKK07200 and BKK07210, respectively) (**Fig. 3B**, Δ*yetK* shown in **Fig. S2**). During integration of the *kan^R^* antibiotic cassettes, recombination occurs at the upstream and downstream regions of *<u>yetJ</u>* and *yetK* genes of the 168 strains (used for the BKK library), resulting in removal of the PGC in MB9_B4.

**Fig. 3.**
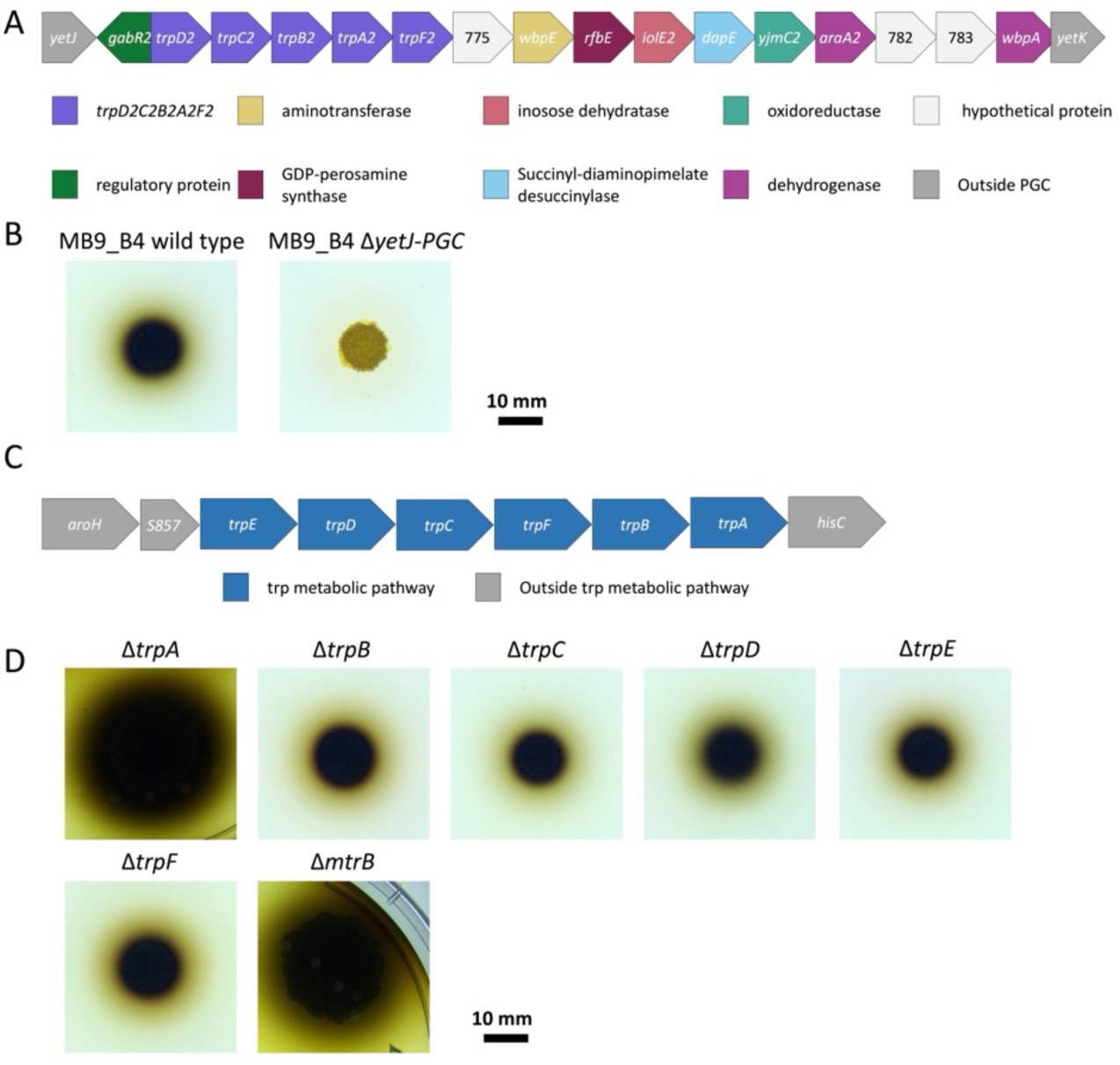
A) Diagram of the PGC, coloration by product class and presumed function from annotation, additional information on each gene can be found in table 1 And 2. B) Deletion of *yetJ*-PGC results in loss of pigmentation on MSgg at 30 °C, image from a 4-day-old samples. C) Diagram of the canonical tryptophan metabolic gene cluster in *B. subtilis* (blue) and surrounding genes for context (grey). D) Pigment cluster is unaffected by deletion of *mtrB* and *trpABCDEF* biosynthetic genes. Scale bar corresponds to 10 mm for the images in B and C.

**Table 1.** Genes in PGC by their abbreviated name used in Fig 3A with name and length annotated by PGAP and bakta.

| Abbreviation | Name (PGAP) | Name (bakta) | Length (PGAP) | Length (bakta) |
| --- | --- | --- | --- | --- |
| 605 | *gabR_4* | JONKOO_20605 | 1416 | 1416 |
| 610/*trpD2* | *trpD_1* | JONKOO_20610 | 1059 | 1059 |
| *trpC2* | *trpC_1* | *trpC* | 783 | 783 |
| *trpB2* | *trpB_1* | *trpB* | 1197 | 1197 |
| *trpA2* | *trpA_1* | *trpA* | 819 | 819 |
| 630 | *trpF_1* | JONKOO_20630 | 654 | 654 |
| 635 | ACGILOGKE_00775 | JONKOO_20635 | 1029 | 486 |
| 640 |  | JONKOO_20640 |  | 474 |
| 645 | *wbpE* | JONKOO_20645 | 1152 | 1152 |
| 650 | *rfbE* | JONKOO_20650 | 1128 | 1128 |
| 655 | *iolE_1* | JONKOO_20655 | 870 | 870 |
| 660 | *dapE* | JONKOO_20660 | 1296 | 1296 |
| 665 | *yjmC_1* | JONKOO_20665 | 1104 | 1104 |
| 670 | *araA_1* | JONKOO_20670 | 903 | 903 |
| *yopX* | ACGILOGKE_00782 | *yopX* | 414 | 414 |
| 680 | ACGILOGKE_00783 | JONKOO_20680 | 690 | 567 |
| 685 | *wbpA* | JONKOO_20685 | 1335 | 1335 |

**Table 2.**
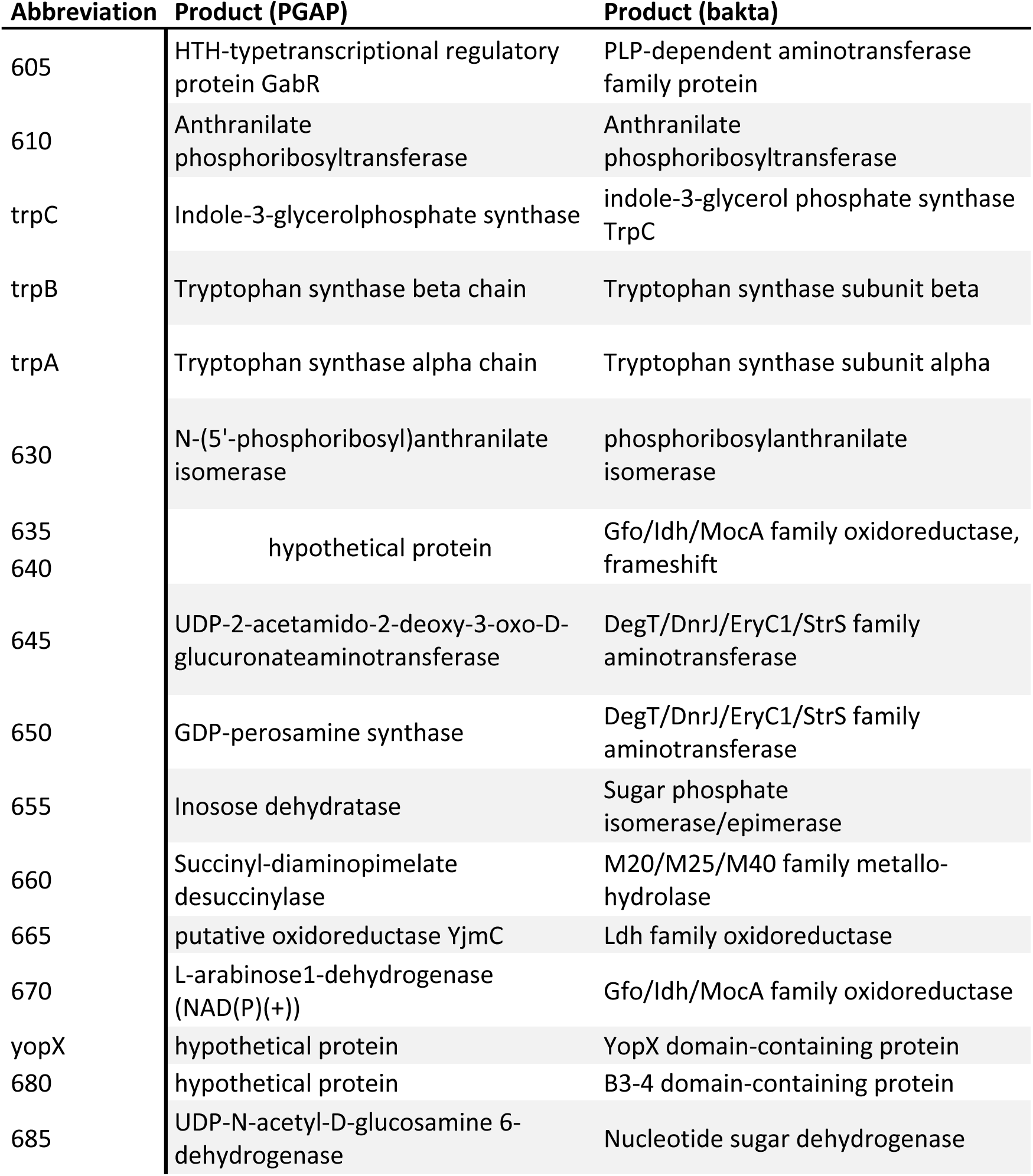
Genes in PGC by their abbreviated name used in Fig 3A with product annotated by PGAP and bakta.

Since multiple genes were annotated to encode part of tryptophan biosynthesis, we next tested if the PGC could complement deletion of the canonical *trpA-F* genes or if pigment production was affected by tryptophan regulatory elements (**Fig. 3C**, other tested mutants can be found in **Fig. S3**). Single deletion mutants of the *trpA-F* genes in MB9_B4 were unable to grow on media lacking tryptophan. Likewise, deletion of the *mtrB* gene encoding the RNA-binding attenuation protein of tryptophan biosynthesis did not affect pigmentation. Thus, these experiment suggests that if PGC genes originated from the canonical *trp* operon, they have been fully repurposed from tryptophan synthesis and regulation beyond the promiscuity that some enzymes exhibit during neofunctionalization (32).

Blast search using the genes of the PGC revealed additional 27 genomes or scaffolds carrying a PGC with high identity (**Fig. 4**). All isolates that carried a PGC belonged to the *B. subtilis* species complex with the majority being either *B. subtilis* or *Bacillus mojavensis*. The PGC itself was remarkably conserved across most PGC-carrying isolates (ANI = 100 – 98.19%) except NRS-0660 which was lacking the first ∼5000 bps and NRS-0276 where only fragments of the PGC could be identified. This might be due to incomplete genome sequences, as the genome sequences of NRS-0660 and NRS-0276 are not completed and closed. Corresponding strains were tested for pigmentation on MSgg agar, here NRS- 0660 was able to produce the pigment while NRS-0276 exhibited a light purple coloration. Out of the remaining 25 isolates, further four were unable to produce pigmentation in the assayed conditions although there were no signs of disruption of the PGC in these.

**Fig. 4.**
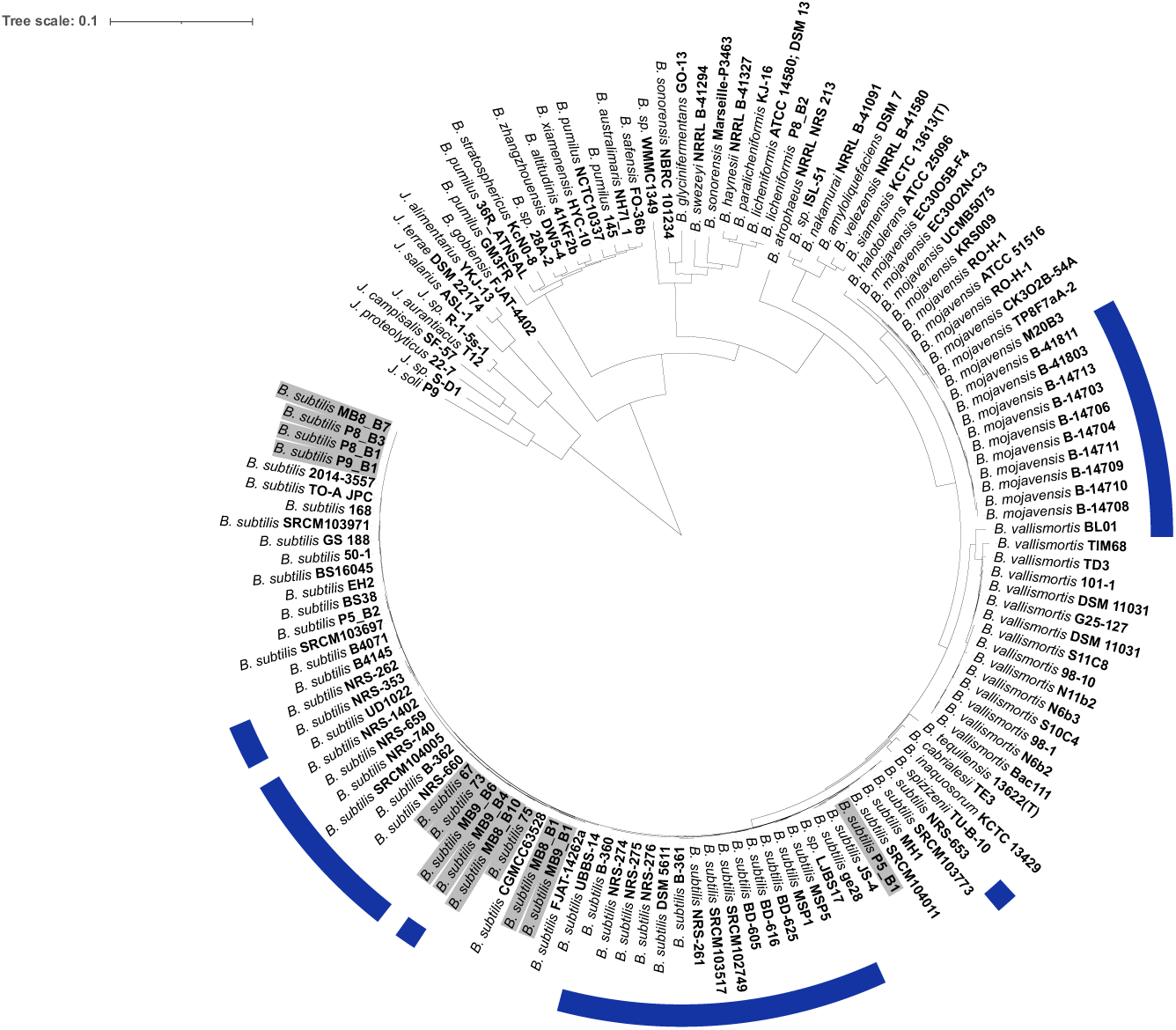
Phylogenetic tree (based on autoMLST) of pigment carrying genomes and closely related strains, PGC-harboring strains are marked with a blue ribbon. Isolates tested in the experiments are highlighted with grey background. Figure created using iTOL (33).

To get more detailed information about the chemical nature of the pigment, we carried out a spectroscopic analysis of *B. subtilis* MB9_B4 and its PGC deleted mutant (**Fig. 5**). Bacterial pellicles constitute extraordinary complex analytical matrices made of many chemical entities (polysaccharides, proteins, amino acids, lipids, water), rendering their high-resolution analysis challenging. Magic-angle spinning solid-state NMR (MAS NMR) and ATR-FTIR spectroscopies constitute powerful analytical tools to investigate the molecular composition of complex insoluble biomolecular pigments (34), as demonstrated for melanin samples of different sources (35, 36). We employed one dimensional ^1^H-^13^C and ^1^H-^15^N insensitive nuclei enhanced by polarization transfer (INEPT) experiments under magic-angle spinning condition (11 kHz) on ^13^C, ^15^N-labeled samples to obtain the molecular fingerprint of Bacillus pellicles. The ^13^C-detected experiment (Fig. 5A) reveals a typical complex spectroscopic signature encoding for protein, lipids, polysaccharides and other molecules (37). By comparing wild type and PGC deleted mutant strains, we detected several spectral areas associated with a substantial reduction of ^13^C signals encoding for alkyl (e.g. methylene) groups at ∼32, 36 and 38 ppm for the PGC deleted strain. Additionally, in the range 86-90 ppm, two major signals were absent for the PGC deleted strain, that could be putatively assigned to -<u>CHOH</u> or -CHOAr. Many microbial pigments are nitrogen-containing molecules, and we probed the structural ^15^N environment with a ^1^H-^15^N INEPT (Fig. 5B). We observed a loss of signal at ^15^N ∼125 ppm, a spectral region that usually encodes nitrogen signal of melanin-like pigments (38, 39). Next, we performed ATR-FTIR to identify the chemical bonds by comparing the fingerprints of the same two samples (Fig. 5C). Noticeable differences were observed for absorbance at 2954 cm^-1^ (aliphatic stretching), 1200 cm^-1^ (phenolic C-OH stretching) and 1046 cm^-1^ (C-H aromatic in plane). A strong contribution at 833 cm^-1^ (out-of-plan bending), often observed in melanin-like samples, was also considerably reduced in the mutant sample. Note that a small absorbance difference (∼20 cm^-1^) is observed in our samples (^13^C-labeled) compared to reported values from the literature obtained on unlabeled samples. Altogether, MAS NMR and ATR-FITR pointed to a pigment containing alkyl, C=C and CHO groups.

**Fig. 5.**
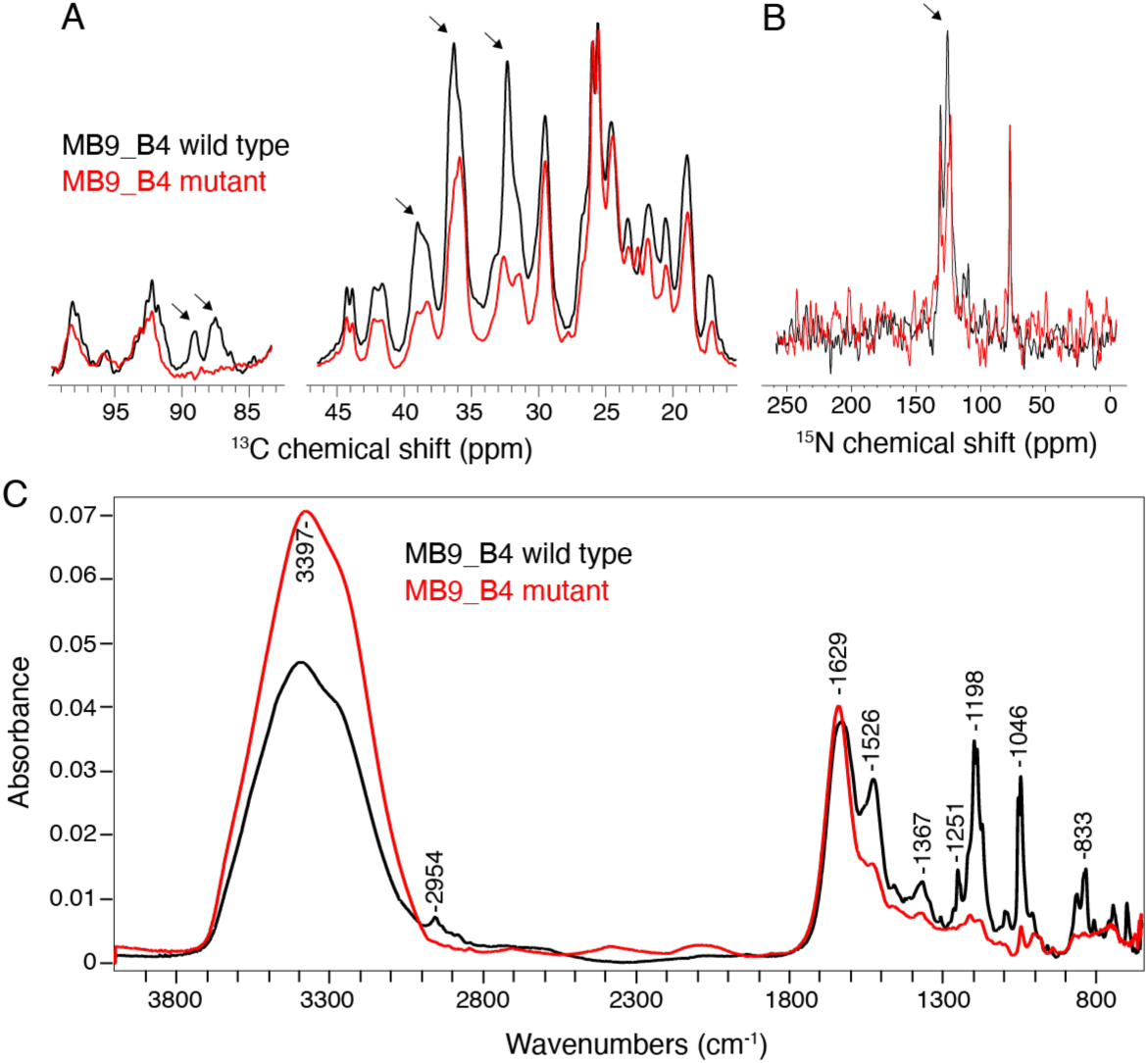
Spectroscopic analysis of ^13^C,^15^N-labeled Bacillus pellicles for the wild type MB9_B4 (black lines) and the mutant lacking PGC (MB9_B4 GFP Δ*yetJ*-PGC, red lines). Magic-angle spinning solid-state NMR analysis of pellicles recorded at 600 MHz at a spinning frequency of 11 kHz, using one dimensional ^1^H- ^13^C and ^1^H-^15^N INEPT spectra, respectively in panel A and B. C) Fourier transform infrared absorbance spectra.

## 4. DISCUSSION

Secondary metabolites provide a huge reservoir of untapped potential, with a large majority of antibiotics and therapeutics originating from microbial products. However, while some classes of SMs are studied in detail, it is hypothesized that a large majority of the biosynthetic potential is still relatively unknown. The current tools for discovery of SMs largely rely on the similarity of the biosynthetic apparatus encoding genes to known gene clusters and are challenged when it comes to finding truly novel SM-related BGCs; thus, examples of novel BGCs serve greatly to increase the available parts of the SM potential.

Our results identify the corresponding BGC for a dark pigment present in certain *Bacillus* species. Such pigment production in *Bacilli* was first observed in 1896 leading to the nomination *B. aterrimus* with later studies focusing on culture conditions and pigment dissociates. The pigment gene cluster (PGC) is, by our knowledge, the first of its class with a high likeness to several primary metabolism genes, which hindered detection by popular BGC mining software tools, including antiSMASH. The PGC identified in this study is a valuable addition to our knowledge, which we hope to improve bioinformatic tools for BGC mining in the future.

The function of the pigment is, as of yet, unknown. However, the pattern of its expression during biofilm development could give some indications towards the pigment being beneficial during sporulation or the biofilm life-style. Pigmentation as a strategy for protecting spores against UV and other environmental stressors has been documented in many spore-formers including *Bacillus* itself; In (40), a carotene-like pigment in the *Bacillus atrophaeus* endospore coat was shown to have a protective effect against UV-A radiation. However, the fact that the pigment in our study easily diffuses into the environment and is not bound to the cell or spore coat, questions whether a diffusible protective compound would be advantageous protecting the spores directly from UV radiation or it might have a more global, population level protective function.

Other water-soluble pigments such as phenazines function as an electron shuttle, which provides subpopulations of the *Pseudomonas aeruginosa* biofilm access to electron acceptors and donors (41).

*B. subtilis* produces highly complex biofilms both on solid surfaces, but more interestingly as a pellicle in which a second sub-population residing in the oxygen-poor media beneath the pellicle exists (42). A similar system could greatly benefit biofilm growth in *B. subtilis* by allowing co-metabolism between certain subpopulations by a diffusible metabolite. However, we note that the pigment examined in our study seemed mainly to react irreversibly with oxidizing agents with no visible change under reductive conditions and it is thus likely a poor candidate as an electron shuttle.

Our spectroscopic analysis by MAS NMR and FTIR identified characteristic chemical groups in the pigments, with methylene, C=C, -CHO-C=C- and C-OH. Several FTIR absorbance peaks would virtually fit with the presence of a melanin-like compound. In addition, MAS NMR reveals the presence of a nitrogen-containing pigment, also present in melanin-like molecules. While our spectroscopic study focused on native pellicles, most studies from the literature have been carried out on purified microbial melanin samples (43–46), therefore several spectral ambiguities hamper the exact chemical structure determination of the pigment. Further, advanced purification protocols will be required to drastically simplify the pellicle sample and obtain pure pigment for high resolution structural analysis.

A previous study on *B. aterrimus* from 1946 described a gradual loss of capability to produce the pigment on potato when serially cultured in glucose broth or on glucose-nitrate agar (1 g/L K_2_HPO_4_, 1g/L NaNO_3_, 13 g/L agar, 10 g/L glucose), which could sometimes be regained by serial culturing on potato again (21). Glucose-nitrate media and potato differ widely in their composition, but such difference could be the key to determine the benefit of pigment production as the growth on potato conveyed advantage for pigment producing derivatives and allowed pigmentation to reappear in the strain that had otherwise lost pigment production. It is notable that other pigments and phenotypes were also described in the original and later works (19–21), which likewise might have been lost due to taxonomic changes in species names and due to the use of different laboratory protocols. Pigmentation provides the evidence of a compound without necessarily knowing either the function or corresponding gene cluster, but lack of documented mechanistic knowledge on the synthesis likely increases the novelty of the gene cluster. Therefore, these visible SMs are a promising resource for improving our SM discovery bioinformatic tools.

*B. subtilis* and several species in the *Bacillus* genus are recognized as GRAS (generally recognized as safe) organism and is already used as an additive or in fermentation in a large range of food products (47). As the pigment is produced in food grade materials such as potato plugs, it might have application as a natural colorant and possible anti-oxidant for foodstuffs; although, further tests should be carried out to ensure the safety of the pigment as animal feed or for human consumption.

## Acknowledgments

This project was supported by the Danish National Research Foundation (DNRF137) for the Center for Microbial Secondary Metabolites and the Novo Nordisk Foundation for the “Imaging microbial language in biocontrol (IMLiB)” infrastructure grant (NNF19OC0055625). This work has benefited from the Biophysical and Structural Chemistry Platform at IECB, CNRS UAR 3033, INSERM US001.

## Supplementary files

**Fig S1.**
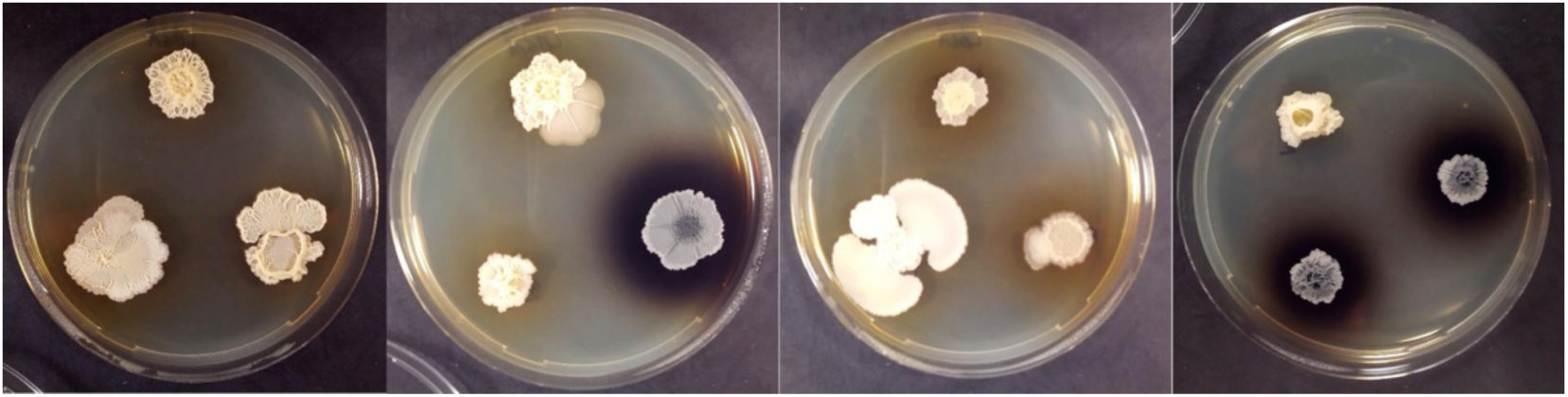
Screening of soil isolates. First plate from the left: MB8_B1 (left), MB8_B7 (right), and MB8_B10 (bottom). Second plate from the left: MB9_B1 (left), MB9_B4 (right), and MB9_B6 (bottom). Third plate from the left: P8_B1 (left), P8_B3 (right), and P9_B1 (bottom). Fourth plate from the left: 73 (left), 67 (right), and P5_B1 (bottom).

**Fig S2.**
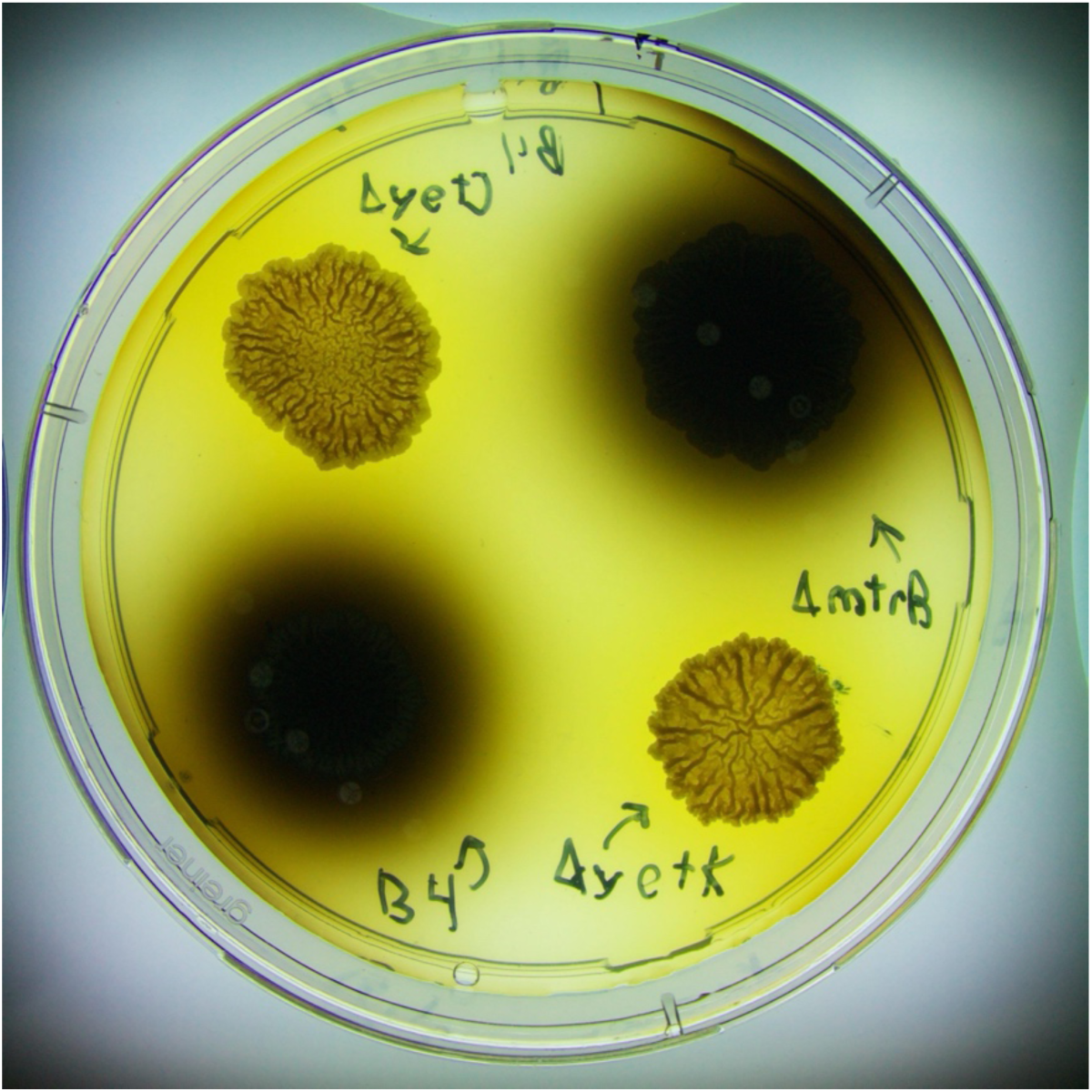
MB9_B4 mutants tested for pigment production. Deletion of PGC accompanied with Δ*yetJ* (top left) or Δ*yetK* (bottom right), deletion of Δ*mtrB* (top right), and wild-type MB9_B4 (bottom left).

**Fig S3.**
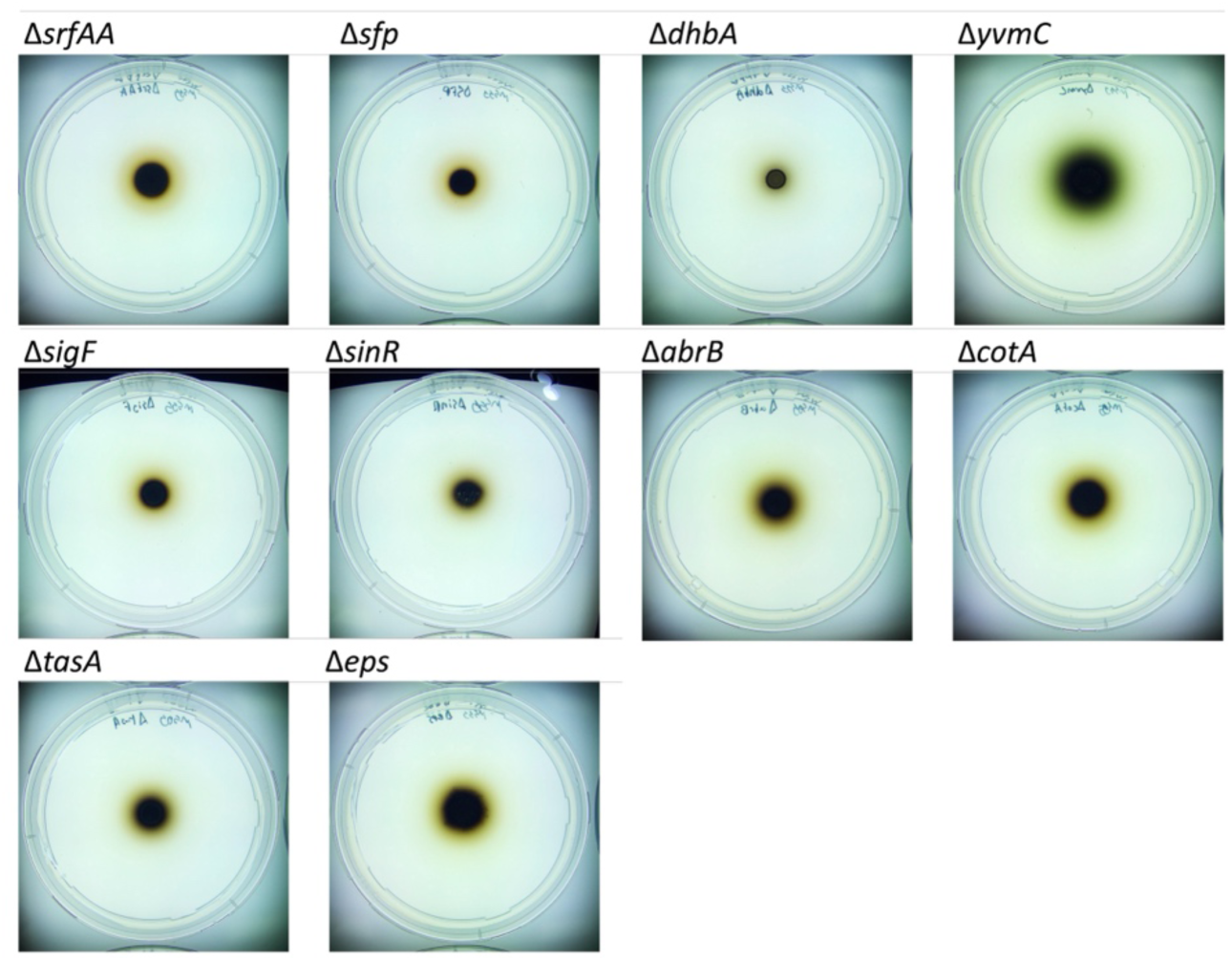
MB9_B4 mutants tested for pigment production.

**Table S1.**
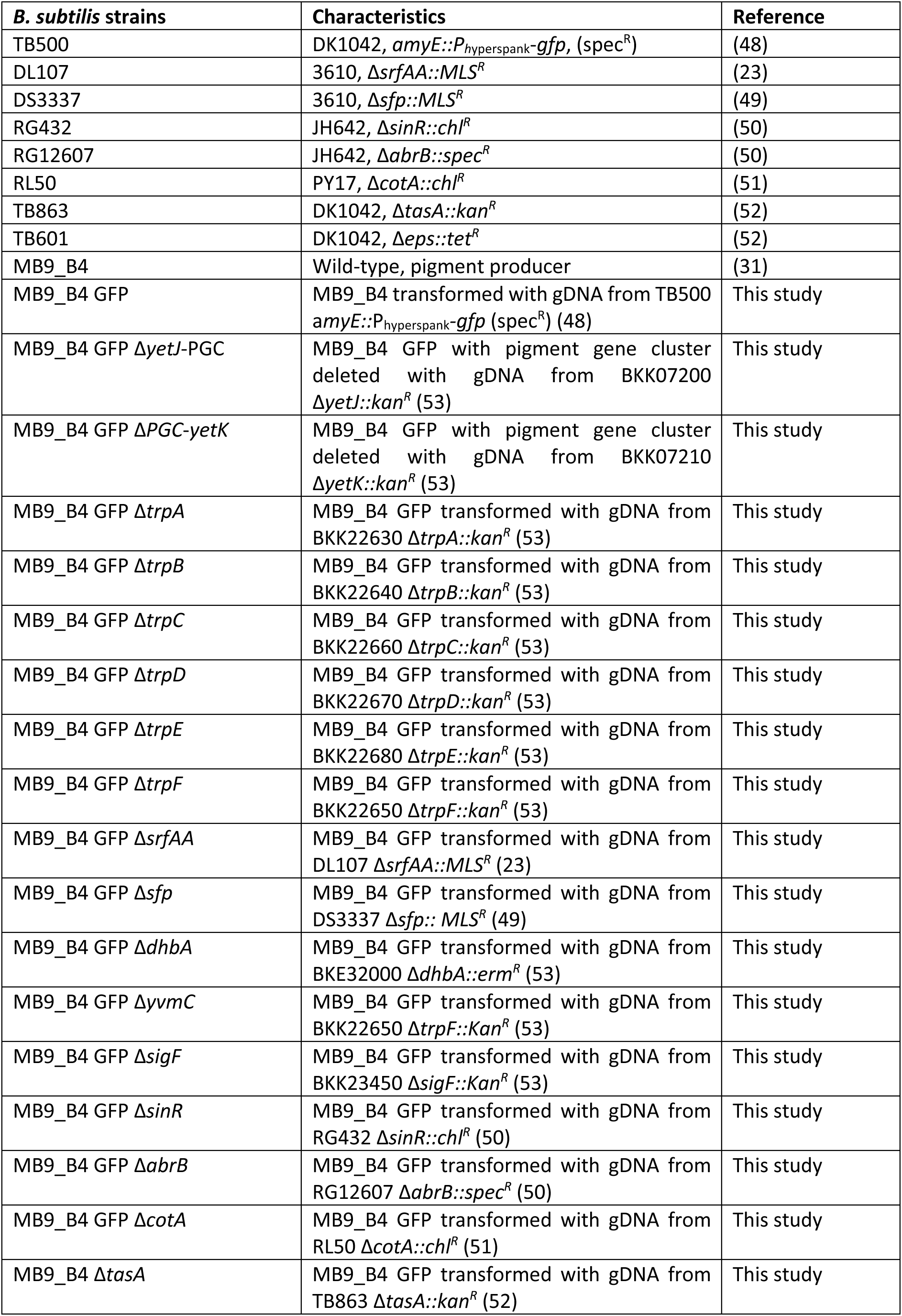
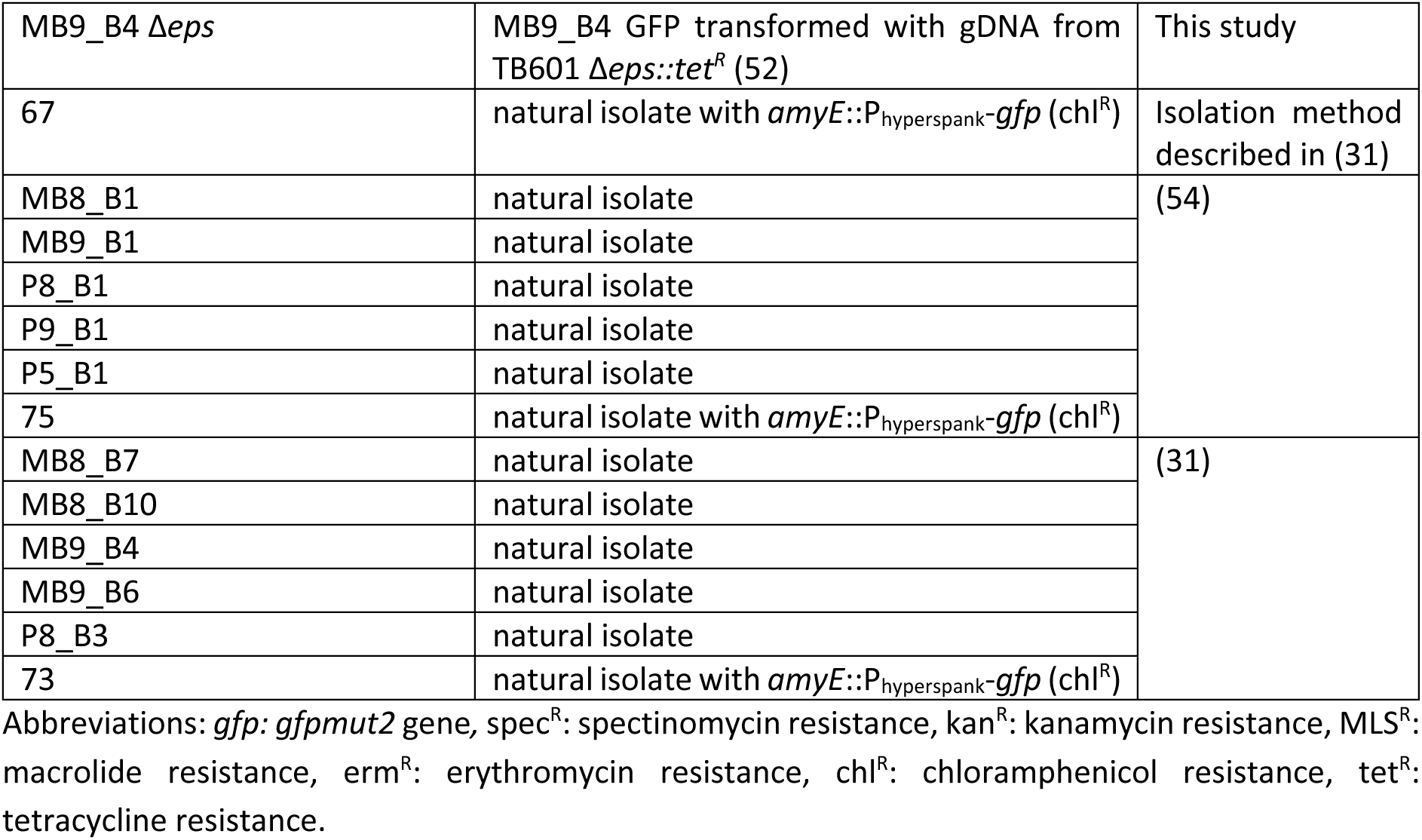
Strains used in this study.

| ***B. subtilis* strains** | **Characteristics** | **Reference** |
| --- | --- | --- |
| TB500 | DK1042, *amyE::P<sub>hyperspank</sub>-gfp*, (*spec<sup>R</sup>*) | (48) |
| DL107 | 3610, *ΔsrfAA::MLS<sup>R</sup>* | (23) |
| DS3337 | 3610, *Δsfp::MLS<sup>R</sup>* | (49) |
| RG432 | JH642, *ΔsinR::chl<sup>R</sup>* | (50) |
| RG12607 | JH642, *ΔabrB::spec<sup>R</sup>* | (50) |
| RL50 | PY17, *ΔcotA::chl<sup>R</sup>* | (51) |
| TB863 | DK1042, *ΔtasA::kan<sup>R</sup>* | (52) |
| TB601 | DK1042, *Δeps::tet<sup>R</sup>* | (52) |
| MB9_B4 | Wild-type, pigment producer | (31) |
| MB9_B4 GFP | MB9_B4 transformed with gDNA from TB500 *amyE::P<sub>hyperspank</sub>-gfp* (*spec<sup>R</sup>*) (48) | This study |
| MB9_B4 GFP *ΔyetJ*-PGC | MB9_B4 GFP with pigment gene cluster deleted with gDNA from BKK07200 *ΔyetJ::kan<sup>R</sup>* (53) | This study |
| MB9_B4 GFP *ΔPGC-yetK* | MB9_B4 GFP with pigment gene cluster deleted with gDNA from BKK07210 *ΔyetK::kan<sup>R</sup>* (53) | This study |
| MB9_B4 GFP *ΔtrpA* | MB9_B4 GFP transformed with gDNA from BKK22630 *ΔtrpA::kan<sup>R</sup>* (53) | This study |
| MB9_B4 GFP *ΔtrpB* | MB9_B4 GFP transformed with gDNA from BKK22640 *ΔtrpB::kan<sup>R</sup>* (53) | This study |
| MB9_B4 GFP *ΔtrpC* | MB9_B4 GFP transformed with gDNA from BKK22660 *ΔtrpC::kan<sup>R</sup>* (53) | This study |
| MB9_B4 GFP *ΔtrpD* | MB9_B4 GFP transformed with gDNA from BKK22670 *ΔtrpD::kan<sup>R</sup>* (53) | This study |
| MB9_B4 GFP *ΔtrpE* | MB9_B4 GFP transformed with gDNA from BKK22680 *ΔtrpE::kan<sup>R</sup>* (53) | This study |
| MB9_B4 GFP *ΔtrpF* | MB9_B4 GFP transformed with gDNA from BKK22650 *ΔtrpF::kan<sup>R</sup>* (53) | This study |
| MB9_B4 GFP *ΔsrfAA* | MB9_B4 GFP transformed with gDNA from DL107 *ΔsrfAA::MLS<sup>R</sup>* (23) | This study |
| MB9_B4 GFP *Δsfp* | MB9_B4 GFP transformed with gDNA from DS3337 *Δsfp::MLS<sup>R</sup>* (49) | This study |
| MB9_B4 GFP *ΔdhbA* | MB9_B4 GFP transformed with gDNA from BKE32000 *ΔdhbA::erm<sup>R</sup>* (53) | This study |
| MB9_B4 GFP *ΔyvmC* | MB9_B4 GFP transformed with gDNA from BKK22650 *ΔtrpF::Kan<sup>R</sup>* (53) | This study |
| MB9_B4 GFP *ΔsigF* | MB9_B4 GFP transformed with gDNA from BKK23450 *ΔsigF::Kan<sup>R</sup>* (53) | This study |
| MB9_B4 GFP *ΔsinR* | MB9_B4 GFP transformed with gDNA from RG432 *ΔsinR::chl<sup>R</sup>* (50) | This study |
| MB9_B4 GFP *ΔabrB* | MB9_B4 GFP transformed with gDNA from RG12607 *ΔabrB::spec<sup>R</sup>* (50) | This study |
| MB9_B4 GFP *ΔcotA* | MB9_B4 GFP transformed with gDNA from RL50 *ΔcotA::chl<sup>R</sup>* (51) | This study |
| MB9_B4 *ΔtasA* | MB9_B4 GFP transformed with gDNA from TB863 *ΔtasA::kan<sup>R</sup>* (52) | This study |
| MB9_B4 $\Delta eps$ | MB9_B4 GFP transformed with gDNA from TB601 $\Delta eps::tet^R$ (52) | This study |
| 67 | natural isolate with *amyE::P<sub>hyperspank</sub>-gfp* (chl<sup>R</sup>) | Isolation method described in (31) |
| MB8_B1 | natural isolate | (54) |
| MB9_B1 | natural isolate |  |
| P8_B1 | natural isolate |  |
| P9_B1 | natural isolate |  |
| P5_B1 | natural isolate |  |
| 75 | natural isolate with *amyE::P<sub>hyperspank</sub>-gfp* (chl<sup>R</sup>) |  |
| MB8_B7 | natural isolate | (31) |
| MB8_B10 | natural isolate |  |
| MB9_B4 | natural isolate |  |
| MB9_B6 | natural isolate |  |
| P8_B3 | natural isolate |  |
| 73 | natural isolate with *amyE::P<sub>hyperspank</sub>-gfp* (chl<sup>R</sup>) |  |

Abbreviations: *gfp*: *gfpmut2* gene, spec<sup>R</sup>: spectinomycin resistance, kan<sup>R</sup>: kanamycin resistance, MLS<sup>R</sup>: macrolide resistance, erm<sup>R</sup>: erythromycin resistance, chl<sup>R</sup>: chloramphenicol resistance, tet<sup>R</sup>: tetracycline resistance.

